# Susceptibility rhythm to bacterial endotoxin in myeloid clock-knockout mice

**DOI:** 10.1101/766519

**Authors:** Veronika Lang, Sebastian Ferencik, Bharath Ananthasubramaniam, Achim Kramer, Bert Maier

## Abstract

Local circadian clocks are active in most cells of our body. However, their impact on circadian physiology is still under debate. Mortality by endotoxic (LPS) shock is highly time-of-day dependent and local circadian immune function such as the cytokine burst after LPS challenge has been assumed to be causal for the large differences in survival. Here, we investigate the roles of light and myeloid clocks on mortality by endotoxic shock. Strikingly, mice in constant darkness (DD) show a three-fold increased susceptibility to LPS as compared to mice in light-dark conditions. Mortality by endotoxic shock as a function of circadian time is independent of light-dark cycles as well as myeloid CLOCK or BMAL1 as demonstrated in conditional knockout mice. Unexpectedly, despite the lack of a myeloid clock these mice still show rhythmic patterns of pro- and anti-inflammatory cytokines such as TNF*α*, MCP-1, IL-18 and IL-10 in peripheral blood as well as time-of-day and site dependent traffic of myeloid cells. We speculate that systemic time-cues are sufficient to orchestrate innate immune response to LPS by driving immune functions such as cell trafficking and cytokine expression.

## Introduction

Timing of immune-functions is crucial for initiating, establishing, maintaining and resolving immune-responses. The temporal organization of the immune system also applies for daily recurring tasks and may even help to anticipate times of environmental challenges. In humans, many parameters and functions of the immune system display diurnal patterns [1], which impact on disease severity and symptoms [2]. While the concepts of chronobiology are increasingly acknowledged in life-science and medicine [3], a deep comprehension of how time-of-day modulates our physiology in health and disease is still lacking.

The fundamental system behind the time-of-day dependent regulation of an organism, its behavior, physiology and disease is called the circadian clock. In mammals, this clock is organized in a hierarchical manner: a central pacemaker in the brain synchronized to environmental light-dark cycles via the eyes and peripheral clocks receiving and integrating central as well as peripheral (e.g. metabolic) time information. Both central and peripheral clocks are essentially identical in their molecular makeup: Core transcription factors form a negative feedback loop consisting of the activators, CLOCK and BMAL1, and the repressors, PERs and CRYs. Additional feedback loops and regulatory factors amplify, stabilize and fine-tune the cell intrinsic molecular oscillator to achieve an about 24-hour (circadian) periodicity of cell- and tissue-specific clock output functions [4].

Circadian patterns of various immune-functions have been reported in mice [1] and other species [5–7] including cytokine response to bacterial endotoxin and pathogens, white blood cell traffic [8–10] and natural killer cell activity [11]. Cell-intrinsic clocks have been described for many leukocyte subsets of lymphoid as well as myeloid origin, including monocytes/macrophages [12]. Furthermore, immune-cell intrinsic clocks have been connected to cell-type specific output function such as the TNF*α* response to LPS in macrophages [12].

Sepsis is a severe life threatening condition with more than 31 million incidences per year worldwide [13]. Mouse models of sepsis show a strong time-of-day dependency in mortality rate when challenged at different times of the day [14–18]. Most studies agree about the times of highest (around *Zeitgeber time* [ZT]8 - i.e. 8 hours past lights on) or lowest (around ZT20) mortality across different animal models and investigators [14–17]. The fatal cascade in the pathomechanism of endotoxic shock is initiated by critical doses of LPS recognized by CD14 bearing monocytes/macrophages leading to a burst of pro-inflammatory cytokines. A viscous cycle of leukocyte recruitment, activation and tissue factor (III) expression triggers disseminated intravascular coagulation and blood pressure decompensation and final multi-organ dysfunction [19].

Here, we investigate the impact of light-dark cycles and local myeloid clocks on time-of-day dependent survival rates in endotoxic shock using conditional clock-knockout mouse models. We show that peripheral blood cytokine levels as well as mortality triggered by bacterial endotoxin depend on time-of-day despite a functional clock knockout in myeloid cells. Our work thus challenges current models of local regulation of immune responses.

## Results

### Time-of-day dependent survival in endotoxic shock

Diurnal patterns of LPS-induced mortality (endotoxic shock) have been reported numerous times in different laboratory mouse strains [9, 14, 20]. To address the question, whether time-of-day dependent susceptibility to LPS is under control of the circadian system rather than being directly or indirectly driven by light, we challenged mice kept either under light-dark conditions (LD 12:12) or in constant darkness (DD) at four different times during the cycles. As expected from previous reports, survival of mice housed in LD was dependent on the time of LPS injection, being highest during the light phase and lowest at night (Fig. 1A, Suppl. Fig. 1A-C).

**Figure 1.**
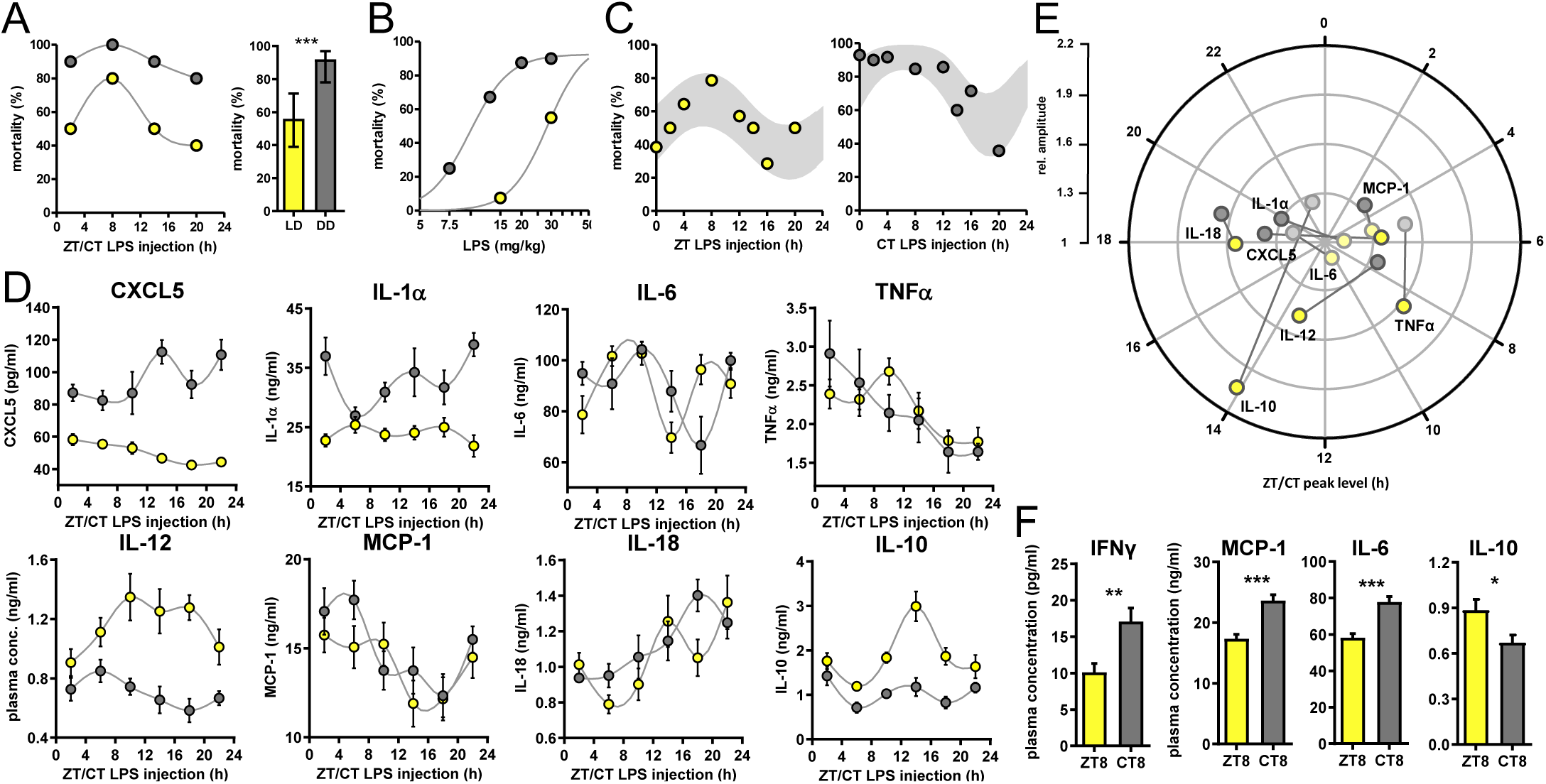
Time-of-day dependent mortality in LPS treated mice is controlled by the circadian system and light conditions. **A)** LPS (30mg/kg, *i*.*p*.) induced mortality in C57Bl/6 mice (n=10 per time point) kept either in LD 12:12 (yellow) or in DD (grey). Left graph: single time points; right graph: mean mortality of LD or DD light condition. Error bars represent 95% confidence intervals (n=40 per group, *** p<0.001). **B)** LPS dose-mortality curves of mice challenged at 4 time points in LD versus DD (overall n=40 mice per dose). Gray lines were calculated by fitting an allosteric model to each group. **C)** Mice (n=10-14 per time point) were challenged with half-lethal doses of LPS (30 mg/kg, *i*.*p*., for mice kept in LD (left panel) or 13mg/kg, *i*.*p*., for mice kept in DD (right panel). Mortality was assessed 60 hours after LPS injection. To perform statistical analyses, mortality rates were transformed to probability of death in order to compute sine fit using logistic regression and F-test (LD, p=0.06; DD, p=0.001; gray shaded areas indicate 95% confidence intervals. **D) and E)** Time-of-day dependent cytokine profiles in peripheral blood of C57Bl/6 mice (n=10 per time point) challenged with half-lethal doses of LPS (30mg/kg, *i*.*p*., for mice kept in LD (yellow) or 13mg/kg, *i*.*p*., for mice kept in DD (gray)) and sacrificed 2 hours later. **E**) Relative amplitudes and phases of cytokines shown in D). Light-colored circles represent non-significant circadian rhythms (p-value>0.05) as determined by non-linear least square fit and consecutive F-test (see also Methods section). **F)** Cytokine levels in peripheral blood from mice (n=10 per time point) challenged with LPS (13mg/kg, *i*.*p*.) at either ZT8 (LD) or CT8 (DD) conditions (T-test, *** p<0.001, ** p<0.01, *p<0.05).

Surprisingly, mice challenged one day after transfer in constant darkness using the same dose showed a more than 60% increase in overall mortality compared to mice kept in LD (Fig. 1A). Furthermore, time-of-day dependent differences in mortality were much less pronounced under these conditions.

To discriminate, whether constant darkness alters overall susceptibility to LPS leading to a ceiling effect rather than eliminating time-dependent effects, we systematically reduced LPS dosage in DD conditions. By challenging the mice at four times across the day we determined the half-lethal dose of LPS in constant darkness (Fig. 1B). As suspected, mice in DD showed a 3-fold increased susceptibility to LPS-induced mortality. For circadian rhythm analysis, we extended the number of time points at approximately half-lethal doses both in LD and DD groups, respectively. This revealed diurnal/circadian patterns in mortality rate in LD (p-value=0.06) as well as in DD conditions (p-value=0.001) (Fig. 1C) demonstrating that the circadian system controls susceptibility to LPS. In addition, this susceptibility is overall increased in constant darkness (Fig. 1A,B and Suppl. Fig. 1D).

### Circadian cytokine response upon LPS challenge in mice

We and others have recently found that cells in the immune system harbor self-sustained circadian oscillators, which shape immune functions in a circadian manner [12, 21]. Pro-inflammatory responses in murine *ex vivo* macrophage culture [12, 22] are controlled by cell-intrinsic clocks and are most prominent during the day and lowest during the activity phase. Furthermore, pro-inflammatory cytokines such as TNF*α*, IL-1*α*/*β*, IL-6, IL-18 as well as MCP-1 have been linked to the pathomechanism of endotoxic shock [23–27].

Thus, we hypothesized that the cytokine response in LPS-challenged mice has a time-of-day dependent profile, which might govern mortality rate in the endotoxic shock model. Indeed, plasma of mice collected two hours after administration of half-lethal doses of LPS (either in LD (30mg/kg) or DD (13mg/kg) conditions at various times during the day), exhibited an up to two-fold time-of-day difference in absolute cytokine concentrations (Fig. 1D and Suppl. Fig. 1E; Fig. 1E and Suppl. Fig. 1F for amplitude and phase information as determined by sine fit). In animals kept in LD, TNF*α*, showed highest levels around ZT8, IL-18 levels peaked at ZT18 and IL-12 as well as the anti-inflammatory cytokine IL-10 had their peak-time around ZT14. Interestingly, cytokine profiles from DD mice differed substantially from LD profiles: the peak-times of IL-12 was phase-advanced by 6 hours, CXCL5 completely reversed its phase, whereas IL-18 remained expressed predominantly in the night. Taken together, cytokine profiles of LPS-challenged mice parallel endotoxic shock-induced mortality patterns, although most circadian cytokines show variable phase relations between free-running and entrained conditions (Fig. 1E).

Next, we investigated, whether the increased overall mortality in DD was correlated with an increased pro-inflammatory cytokine response. We thus injected mice at either ZT8 (mice kept in LD) or CT8 (mice kept in DD) with the same dose of LPS (13mg/kg) and took blood samples two hours later. IFN*γ*, MCP-1, IL-6 and the anti-inflammatory cytokine IL-10 showed significantly altered levels between mice kept in LD or DD (Fig. 1F), suggesting that in DD conditions the sensitivity to endotoxin is increased leading to enhanced pro-inflammatory cytokine secretion and subsequently increased mortality.

### Dispensable role of myeloid clocks in circadian endotoxin reactivity

Local clocks are thought to play important roles in mediating circadian modulation of cell- and tissue-specific functions [28]. In fact, depletion of local immune clocks has been shown to disrupt circadian patterns of tissue function [29–33]. To test whether local clocks in cells of the innate immune system are responsible for circadian time dependency in the response to bacterial endotoxin, we challenged myeloid lineage *Bmal1*-knockout mice (*LysM-Cre*^+/+^ x *Bmal1*^flox/flox^, hereafter called myBmal-KO) with half-lethal doses of LPS at various times across the circadian cycle. These mice lack physiological levels of Bmal1 mRNA and protein in cells of myeloid origin (Suppl. Fig. 2A-C) and have been characterized in more detail elsewhere [31]. To our surprise, mortality of these mice was still dependent on circadian time of LPS administration (Fig. 2A) indicating that a functional circadian clock in myeloid cells is not required for time-of-day dependent LPS-sensitivity. However, the overall susceptibility to LPS decreased two-fold compared to wild-type mice (Fig. 2B-D and Suppl. Fig. 2D) suggesting that BMAL1 levels in myeloid cells directly or indirectly modulate susceptibility towards LPS. The latter effect could only in part be attributed to genetic background (note decreased susceptibility to LPS in *LysM-Cre*^+/+^ control mice (Fig. 2C and Suppl. Fig. 2D) together arguing for a tonic rather than temporal role of myeloid BMAL1 in regulating LPS sensitivity.

**Figure 2.**
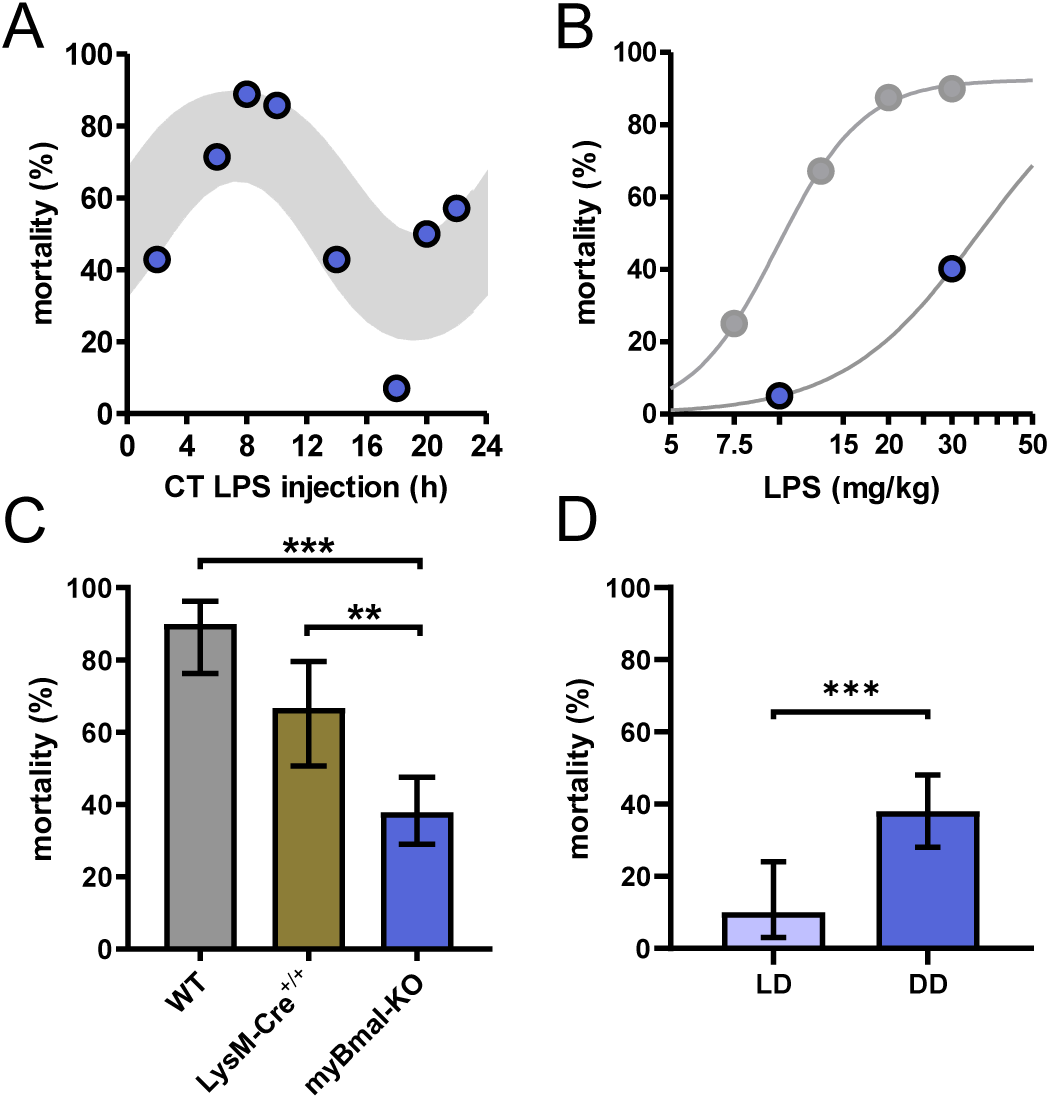
myBmal-KO mice show decreased and circadian time dependent susceptibility to LPS. **A)** Circadian mortality in myBmal-KO mice. Mice kept in DD (n=10-14 per group) were challenged with half-lethal doses of LPS (30mg/kg, *i*.*p*.) at indicated time points. Mortality was assessed 60 hours after LPS injection. Statistics were performed as in Fig. 1*C*, (p=0.0009; gray shaded area indicates 95% confidence interval). **B)** LPS dose-mortality curves of mice challenged at 4-8 time points accross 24 hours in constant dark conditions. About 3-fold decrease of susceptibility to LPS in myBmal-KO mice (blue circles) as compared to wild-type mice (gray circles – replotted from Fig. 1B) kept in DD. Gray lines were calculated by fitting an allosteric model to each group. **C)** Reduced mean mortality in myBmal-KO mice (n=84) compared to control strains LysM-Cre^+/+^ (n=40) or C57Bl/6 wild-type (WT, n=40). All mice were kept in DD and challenged with 30mg/kg LPS *i*.*p*.. Mean values and 95% confidence intervals from WT and myBmal-KO mice were calculated from experiments shown in Fig. 1C (WT) and Fig. 2A (myBmal-KO). **D)** Constant dark conditions render mice more susceptible to LPS independent of Bmal1 in myeloid lineage cells. (*** p<0.001, ** p<0.01).

Despite its essential role for circadian clock function, BMAL1 has been linked with a number of other non-rhythmic processes such as adipogenesis [34], sleep regulation [35] and cartilage homeostasis [36]. Thus, we asked whether the decreased susceptibility to LPS observed in myBmal-KO mice was due to non-temporal functions of BMAL1 rather than to the disruption of local myeloid clocks. CLOCK, like its heterodimeric binding partner BMAL1, has also been shown to be an indispensable factor for peripheral clock function [37]. If non-temporal outputs of myeloid clocks rather than gene specific functions of myeloid BMAL1 controls the susceptibility to LPS, a depletion of CLOCK should copy the myBmal-KO phenotype. We therefore generated conditional, myeloid lineage specific Clock-KO mice (hereafter called myClock-KO). As in myBmal1-KO mice, the expression of Cre recombinase is driven by a myeloid specific promoter (LysM) consequently leading to excision of LoxP flanked exon 5 and 6 of the clock gene [38]. myClock-KO mice showed normal locomotor activity levels and circadian rhythm periods as compared to control mice (Suppl. Fig. 3A,B). As expected, the expression of *Clock* mRNA and protein was substantially reduced in peritoneal cavity cells but was normal in liver (Fig. 3A,B).

**Figure 3.**
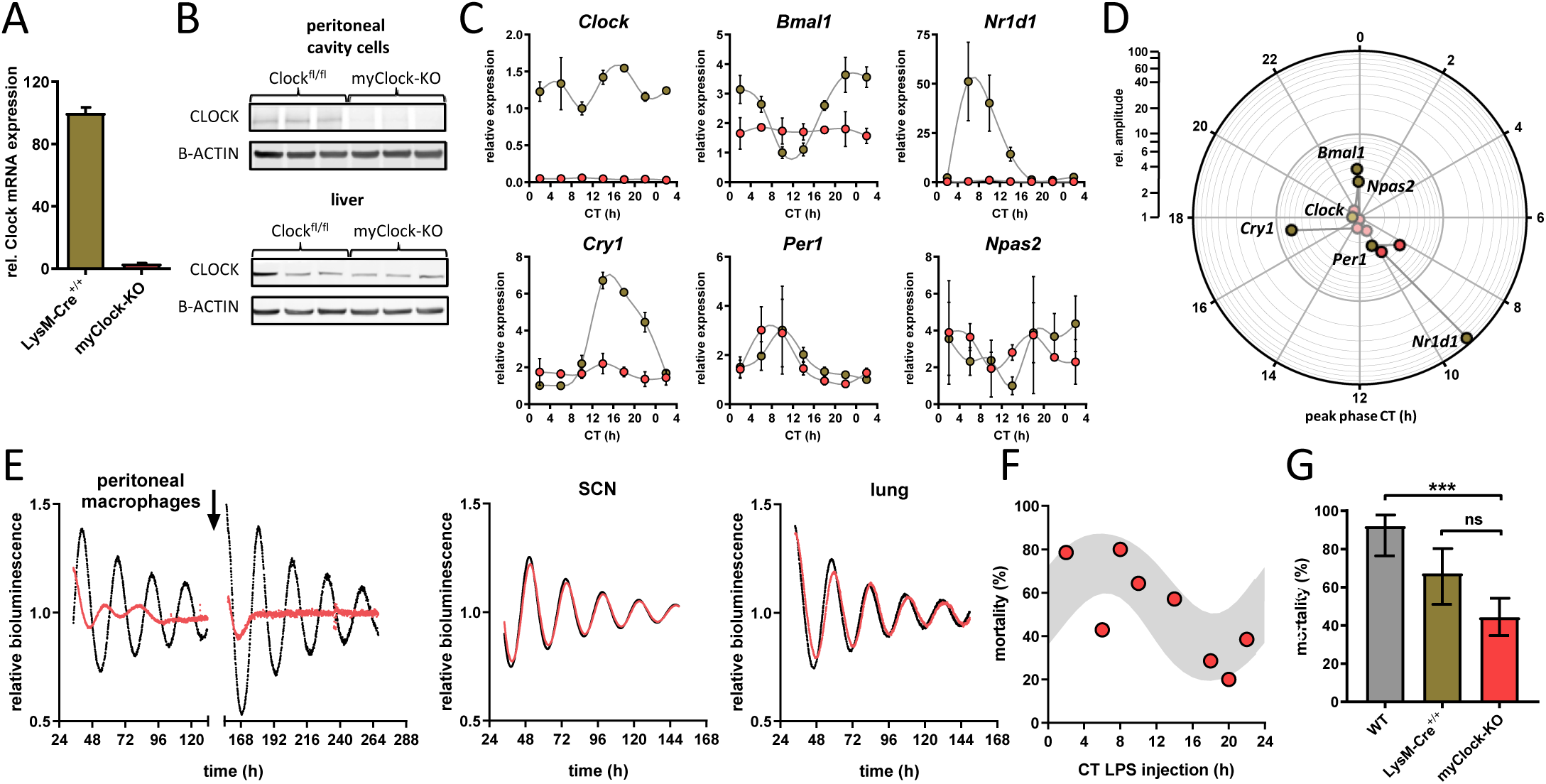
Conditional myClock-KO mice show circadian pattern in mortality by endotoxic shock. **A) and B)** Reduced levels of *Clock* mRNA and protein in myeloid lineage cells of myClock-KO mice. **A)** Mean values of normalized mRNA expression values of time-series shown in C) (n=21 mice per condition, p<0.0001, t-test). **B)** Protein levels by immunoblot in peritoneal cavity cells and liver of myClock-KO and *Clock*^fl/fl^ control mice (n=3). **C)** Relative mRNA levels of selected clock genes in peritoneal macrophages from LysM-cre^+/+^ (brown circles) or myClock-KO (red circles) mice at indicated circadian times. Phase and amplitude information are depicted in D) as analyzed by Chronolyse. Non-significant circadian expressions (p>0.05) are depicted in light red (myClock-KO) or light brown (LysM-cre control). **E)** Representative bioluminescence recordings of peritoneal macrophages, SCN or lung tissue from myClock-KO or wild-type mice crossed with PER2:Luc reporter mice (color coding as before). Black arrow indicates time of re-synchronization by dexamethason treatment (detrended data). **F)** Circadian pattern in endotoxic shock mortality despite deficiency of CLOCK in myeloid lineage cells. Mice (n=10-14 per time point) were challenged with half-lethal doses of LPS (30mg/kg, *i*.*p*.) at indicated time points. Mortality was assessed 60 hours after LPS injection. Statistic was performed as in Fig. 1C (p=0.005, gray shaded area indicates 95% confidence interval). **G)** Reduced mean mortality (at 30mg/kg LPS) in mice deficient of myeloid CLOCK (n=84) compared to control strains LysM-Cre^+/+^ (n=40) or C57Bl/6 (wild-type, n=40). First two bars where re-plotted from Fig. 2C. Error bars represent 95% confidence intervals (n=40 per group, ns p>0.05, p*** p<0.001.

To investigate, whether clock gene rhythms were truly abolished in myClock-KO mice, we harvested peritoneal macrophages from myClock-KO or control mice (LysM-Cre) in regular 4-hour intervals over the course of 24 hours. Rhythmicity of *Bmal1, Cry1, Cry2, Dbp, Npas2* and *Nr1d1* mRNA levels was essentially eliminated, while a low amplitude rhythmicity was *detected for *Per1* and *Per2* mRNA (Fig. 3C,D and Suppl. Fig. 3C,D). Moreover, circadian oscillations were disrupted in peritoneal cavity cells (mainly macrophages and B-cells) from myClock-KO mice, but not in tissue explants from SCN or lung (Fig. 3E).

Given the disruption of rhythmicity in myClock-KO myeloid cells, we asked whether CLOCK in myeloid cells is required for time-of-day dependent mortality in endotoxic shock. To this end, we challenged myClock-KO mice at various times during the circadian cycle with half-lethal doses of LPS. Again, mortality in these mice was significantly time-of-day dependent (Fig. 3F). As in myBmal1-KO mice, myClock-KO mice showed strongly reduced susceptibility to LPS compared to wild-type mice (Fig. 3G and Suppl. Fig. 3E). Taken together, our data unequivocally show that myeloid clockwork are dispensable for the time-of-day dependency in endotoxic shock. In addition, decreased overall susceptibility suggest a non-temporal, sensitizing role of myeloid CLOCK/BMAL1 in the regulation of endotoxic shock.

### Circadian cytokine response in myClock-KO mice

Our initial hypothesis was built on the assumption that a time-of-day dependent cytokine response determines the outcome in endotoxic shock. Previous results from us and others [12, 21, 31] suggested that local myeloid clocks govern the timing of the pro-inflammatory cytokine response. However, circadian mortality profiles in LPS-challenged myeloid clock-knockout mice (Fig. 2A and 3F) led us to question this model: The circadian cytokine response in plasma is either independent of a myeloid clock or the circadian mortality by endotoxic shock does not require a circadian cytokine response.

To test these mutually not exclusive possibilities, we challenged myClock-KO mice with half-lethal doses of LPS in regular 4-hour intervals over the course of one day. Unexpectedly, cytokine levels in plasma, collected two hours after LPS administration still exhibited circadian patterns for TNF*α*, IL-18, IL-10 (Fig. 4A) (p-values=0.046, 0.001 and 0.009, respectively), very similar to those observed in wild-type animals (Fig. 4B). Other cytokines remained below statistical significance threshold for circadian rhythmicity tests (IL-1*α*) or displayed large trends (IL-6) within this period of time (see also Suppl. Fig. 4A,B). These data suggest that an LPS-induced circadian cytokine response does not depend on a functional circadian clock in cells of myeloid origin. Thus, circadian cytokine expression might still be responsible for time-of-day dependent mortality in endotoxic shock.

**Figure 4.**
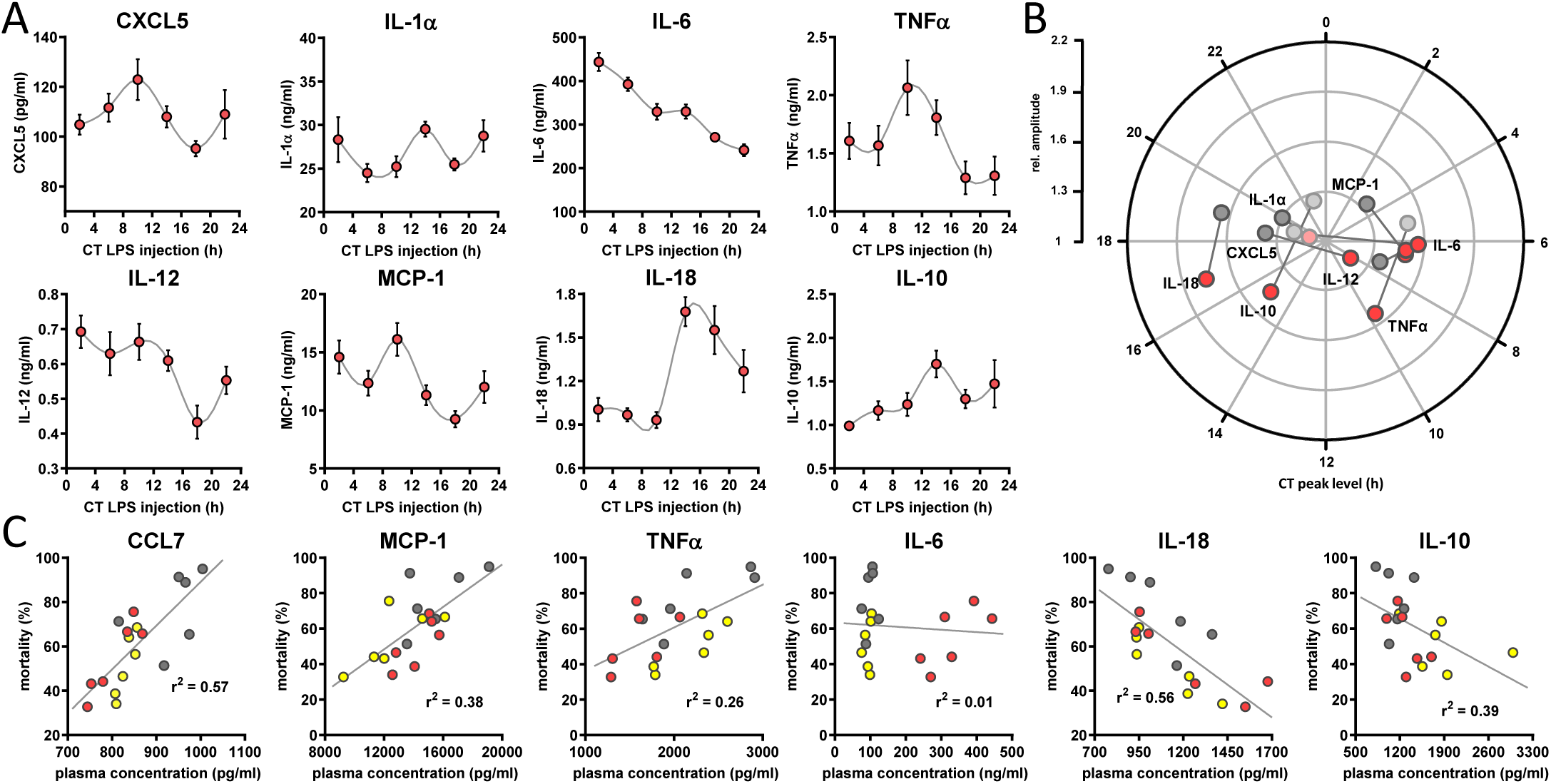
Circadian time dependent cytokine levels in plasma of myClock-KO mice. **A)** Plasma cytokine levels in myClock-KO mice kept in constant darkness, two hours after *i*.*p*. injection of 30mg/kg LPS. Data represent mean values *±* SEM (n=14 per time point). **B)** Polar plot showing amplitude and phase distribution of pro- and anti-inflammatory cytokines from A), red circles and wild-type DD (Fig. 1E). Light circles indicate non-significant circadian abundance (p-values>0.05, non-linear least square fit statistics by Chronolyse). **C)** Overall correlation of cytokine levels with mortality independent of time-of-day of LPS injection and mouse model. Colors indicate data source (wild-type, LD - yellow; wild-type, DD - gray; myClock-KO, DD - red; linear regression - gray line; statistics: spearman correlation).

To identify those cytokines, whose levels best explain mortality upon LPS challenge, we correlated cytokine levels and mortality rate for all conditions - independent of time-of-day or mouse strain - in a linear correlation analysis. Levels of CCL7, MCP-1 and TNF*α* showed strong positive correlation with mortality (p-values=0.0003, 0.0065, 0.0319, respectively). Others, such as IL-10 and IL-18 correlated negatively (p-values=0.0054 and 0.0004, respectively) (Fig. 4C and Suppl. Fig. 4C). Interestingly, while TNF*α* and IL-10 are well known factors in the pathomechanism of endotoxic shock, CCL7, MCP-1 and protective effects of IL-18 have not been reported in this context.

Together, our data suggest that local circadian clocks in myeloid lineage cells are dispensable for time-of-day dependent plasma cytokine levels upon LPS challenge. However, where does time-of-day dependency in endotoxic shock originate instead?

### Persistent circadian traffic in myeloid clock-knockout mice

Circadian patterns in immune cell trafficking and distribution have been recently reported and linked to disease models and immune functions [12, 29, 33]. Similarly, homing and release/egress of hematopoetic stem cells (HSPCs), granulocytes, and lymphocytes to bone marrow and lymph nodes, respectively, have been shown to vary in a time-of-day dependent manner requiring the integrity of an immune-cell intrinsic circadian clock [8–10, 33]. Thus, we asked, whether circadian traffic of myeloid cells can be associated with mortality rhythms in our endotoxic shock model.

To test this, we measured the number of immune cells in various immunological compartments at two distinct circadian time points representing peak (CT8) and trough (CT20) of circadian mortality rate upon LPS challenge. Cells from wild-type, genotype control and myeloid clock-knockout mice (n=5 per time point) were collected from a broad spectrum of immune system compartments (blood, peritoneal cavity, bone marrow, spleen, inguinal lymph nodes, thymus)(Fig. 5A). As expected from previous studies [9, 12, 33] the number of cells were significantly time-of-day dependent in bone marrow, spleen and inguinal lymph nodes of wild-type mice (Fig. 5B, Suppl. Fig. 5B).

**Figure 5.**
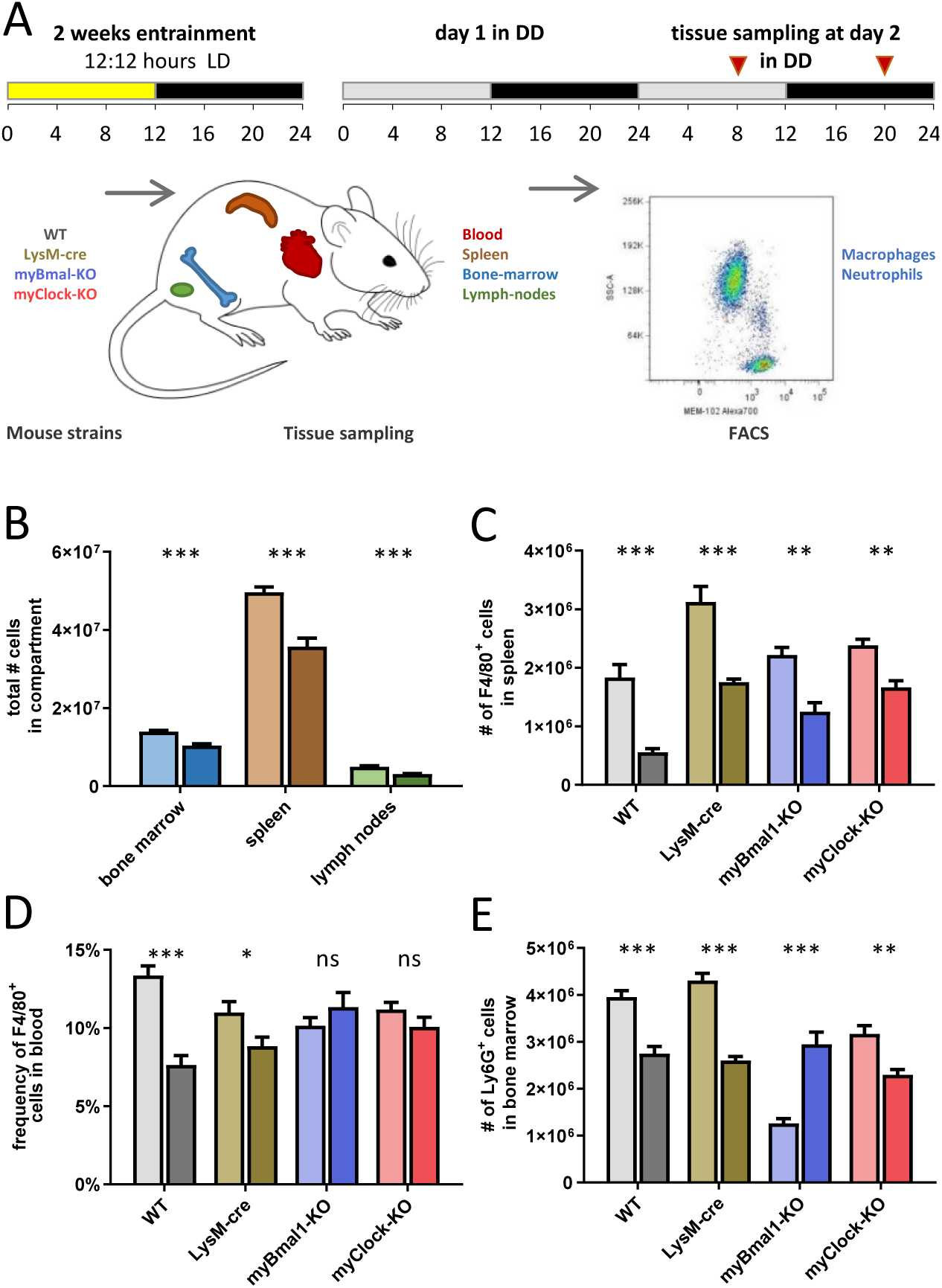
Time of day dependent traffic of myeloid cells despite depletion of myeloid CLOCK or BMAL1. **A)** Experimental scheme to investigate time-of-day dependent immune cell traffic in various compartments and genetic mouse models. **B)** Total cell counts of femoral bone marrow (blue), spleen (brown) or inguinal lymph nodes (green) at CT8 (light colors) or CT20 (dark colors). **C-E)** Cell number or frequency from wild-type and various conditional circadian clock mice at two different circadian time points (wild-type - gray, LysM-cre - brown, myBmal-KO - blue, myClock-KO - red, CT8 - light, CT20 - dark). **C)** Total number of F4/80^+^ macrophages in spleen. **D)** Relative number of F4/80^+^ macrophages in blood. Asterisks indicate level of significance as determined by t-test. **E)** Total number of Ly6G^+^ neutrophils in bone marrow (significance levels: ns p>0.05, * p<0.05, ** p<0.01, *** p<0.001).

If circadian traffic of myeloid cells and therefore lymphoid organ composition would be a main factor in regulating sensitivity to bacterial endotoxin, similar patterns in myeloid clock-knockout mice should be observed. However, results obtained from myBmal-KO and myClock-KO mice did not support this hypothesis. While in spleen, F4/80^+^ macrophages of both myeloid clock knockout strains were still found at higher numbers at CT8 compared to CT20 (Fig. 5C), significant time-of-day differences of monocyte/macrophage numbers in wild-type blood diminished in myeloid clock knockouts (Fig. 5D). Different patterns of time-of-day dependency were also observed in bone marrow (Fig. 5E), together suggesting a rather complex than mono-causal relation between myeloid clocks, circadian traffic and mortality risk in endotoxic shock (for a comprehensive analysis of this dataset see Suppl. Fig. 5B-E).

## Discussion

In this study we tested the hypothesis that local myeloid circadian clocks regulate the devastating immune response in endotoxic shock. A number of findings by various labs including our own pointed to such a possibility. First, monocytes/macrophages have been identified as important cellular entities relaying the endotoxin (LPS) triggered signal to the immune system and other organ systems by means of massive secretion of pro-inflammatory cytokines [39, 40]. Second, a high amplitude circadian clock in monocytes/macrophages has been shown to control circadian cytokine output *in vitro* and *ex vivo* [12, 22, 30]. Third, a high amplitude time-of-day dependent mortality has been demonstrated in endotoxic shock [14]. Fourth, a number of experiments done at two circadian time points, including a cecal ligation and puncture model [18], demonstrating loss of time-of-day dependency of mortality in myeloid *Bmal1* knockout mice [30] further supported this hypothesis.

On the other hand, some studies suggested other mechanisms, e.g. Marpegan and colleagues reported that mice challenged with LPS at two different time points in constant darkness did not exhibit differences in mortality rates [20], arguing for a light-driven process in regulating time-of-day dependency in endotoxic shock. However, we suspected that this result may be caused by a ceiling effect induced by constant darkness, since mortality rates at both time points were close to 100 percent. Indeed, when we compared susceptibility of mice challenged with a similar dose of LPS in light-dark versus constant darkness, we observed a marked increase of overall mortality. Similarly, depletion/mutation of *Per2* rendered mice insensitive to experimental time dependency in endotoxic shock [41]. Surprisingly, however, we found that depletion of either CLOCK or BMAL1 in myeloid lineage derived cells both did not abolish time-of-day dependency in mortality to endotoxic shock, which led us to reject our initial hypothesis.

Our data also exclude light as a stimulus driving mortality: First, data from Halberg, Marpegan, Scheiermann as well as our own lab [9, 12, 14, 20] suggest that mice are more susceptible to endotoxic shock during the light phase compared to dark phase - whereas switching light schedules from light-dark to constant darkness led to an increase in overall mortality. Second, irrespective of the genetic clock-gene depletion tested, our data unequivocally demonstrate circadian mortality rhythms upon LPS challenge even under DD conditions. Strikingly, peripheral blood cytokine levels showed - though altered - circadian time dependency in the myClock-KO strain.

A burst of cytokines, following a lethal dose of LPS is generally thought to be an indispensable factor causing multi-organ dysfunction and leading to death. However, the contribution of single cytokines has been difficult to tease apart due to complex nature of interconnected feedback systems. By adding time-of-day as an independent variable in a number of different mouse models we were able to correlate individual cytokines’ contribution to mortality in the context of the complex response to LPS. While the pro-inflammatory cytokine TNF*α* was confirmed to act detrimentally, IL-18 surprisingly turned out to likely be protective. This interpretation is not only supported by the negative correlation of IL-18 levels in blood and mortality risk, but also by an anti-phasic oscillation of IL-18 levels paralleling those of the known protective cytokine IL-10.

While circadian regulation of trafficking lymphocytes by cell intrinsic clockworks have been demonstrated to impact the pathophysiology of an autoimmune disease model such as EAE [33], our data on distribution patterns of immune cells at two circadian time points draw a more complex picture. In spleen, absolute numbers of macrophages, but not neutrophils and monocytes, are independent of their local clocks. In contrast, patterns of macrophages and neutrophils in peripheral blood and bone marrow (respectively) change upon local myeloid clock depletion. Thus it appears that myeloid cell traffic is regulated at multiple levels including cell-intrinsic, endothelial and site specific factors. However, our data do not support the hypothesis of myeloid cell distribution being a main factor for time-of-day dependent mortality risk in endotoxic shock. Hence, what remains as the source of these rhythms?

Following the patho-physiology of endotoxic shock on its path from cause to effect, it is important to note that the bio-availability of bacterial endotoxin injected intraperitoneally depends on multiple factors (i.e. pharmaco-kinetics), many of which themselves might underlie circadian regulation. As a consequence, same doses of LPS administered *i*.*p*. at different times-of-day might result in highly diverging concentrations at the site of action.

Along this path, rhythmic feeding behavior driven by the central pacemaker was suggested to alter immune-responses directly or indirectly [42, 43]. Also, nutrition related factors could drive immune cells to respond differently to stimuli independent of local clocks [29, 44].

Finally, the ability of cells and organs to resist all sorts of noxa might as well be regulated in a time-of-day dependent manner. In this case, rather than same doses of toxin leading to time dependent cytokine responses (noxa) resulting in diverging rates of multi organ failure, same amount of noxa would cause time dependent rates of multi organ dysfunction and death. Indeed, work from Hrushesky [17] showed that mice challenged with same doses of TNF*α* at various times throughout the day exhibited time-dependent survival paralleling phenomena in endotoxic shock. However, our data showing fluctuating cytokine levels in myClock-KO mice challenged with LPS in constant darkness (Fig. 4A) question the mechanism of organ vulnerability as the only source of circadian regulation.

While our work suggests that local myeloid clocks do not account for time-of-day dependent mortality in endotoxic shock it unequivocally argues for a strong enhancing effect of myeloid CLOCK and BMAL1 on overall susceptibility, which adds on the effect of light conditions. However, it is important to note that our data do stay in conflict with findings from other labs which rather reported attenuating effects of BMAL1 on inflammation [18, 29, 30] but align well with a report on clock mutant mice [22]. Differences in genetic background of mouse strains, animal facility dependent microbiomes [45] or animal care procedures might account for this but remain unsatisfying explanations.

One of the most striking findings of our work is the large increase in susceptibility to endotoxic shock when mice were housed under DD as compared to LD conditions. This increase was observed in wild-type as well as in myBmal-KO mice, which implies that myeloid BMAL1 is not required for this effect. Interestingly, Carlson and Chiu reported similar effects in a cecal ligation and puncture model in rats upon transfer to LL (constant light) or DD conditions, where they found decreased survival as compared to rats remaining in LD conditions [46]. It is tempting to speculate that rhythmic light conditions, rather than light itself promote survival in endotoxemia. In either case, it will be important to further investigate these phenomena not only in respect to animal housing conditions, which need to be tightly light controlled in immunological, physiological and behavioral experiments but also for their apparent implications on health-care in intensive care units.

## Materials and Methods

### Generation of myeloid clock knockout mice

*Bmal1*^flox/flox^ (Bmal-flox) [47] or *Clock*^flox/flox^ (*Clock*-flox) [38] were bred with LysM-Cre to target *Bmal1* or *Clock* for deletion in the myeloid lineage. Offspring were genotyped to confirm the presence of the loxP sites within *Bmal1* or *Clock* and to determine presence of the Cre recombinase. Upon successful recombination the loxP flanked exon 8 of *Bmal1* or in case of *Clock* the floxed exon 5 and 6 were deleted. *LysM*^Cre/Cre^ x *Clock*^flox/flox^ were crossed onto a *Per2:Luc* [48] background for further characterization in bioluminescence reporter assays. All mice have been genotyped before experiments.

### Endotoxic shock experiments

8-12 weeks old mice were entrained to 12h:12h light-dark cycles for 2 weeks. Dosing of LPS injection was adjusted for individual body weight prior injection. Injection volume did not exceed 10*µ*l/g body weight. For LD experiments, mice were injected *i*.*p*. on day 14. For DD experiments mice were transferred to DD on day 14 and were injected *i*.*p*. on second day in DD. Animals were kept in the respective lighting conditions until the termination of experiment 60h post LPS injection. For all endotoxic shock experiments, human endpoints were applied to determine survival (for definition of human endpoints see respective section).

### Time-of-day dependent mortality experiments

Endotoxic shock mortality experiments were comprised of two parts for each mouse strain and condition tested: First, lethal dose 50 (LD_50_) of LPS was determined by injecting groups of mice (n=10) at four 6-hour spaced time points. The LD_50_ describes the concentration at which approx. 50% of all animals injected (averaged over all injection time points) survive. This ensures most dynamic range for the detection of potential circadian rhythms. Second, experimentally determined LD_50_ of LPS was used to investigate, whether mortality by endotoxic shock was dependent on time-of-day. To this end, mice (n=14 per group) were injected *i*.*p*. at six 4-hour spaced time points to increase statistical power for circadian rhythm analysis (see statistical data analysis section).

### Definition of human endpoints

A scoring system was developed in order to detect irreversibly moribund mice before the occurrence of death by endotoxic shock (Suppl. Table 1). It is based on previous reports by [34,41] and required further refinements according to our experience. Mice in the endotoxic shock experiments were monitored and scored every 2-4 hours for up to 60 hours post LPS injection (Suppl. Fig. 1A). In addition, surface body temperature was measured at the sternum every 12h and body weight was measured every 24 h (Suppl. Fig. 1A). Mice with a score of 0-2 were monitored every 4h. As of a score of 3, the monitoring frequency was increased to every 2h. In addition, softened, moisturized food was provided in each cage as of a score of 3. Weight loss exceeding 20%, 3 consecutive scores of 4, or one score of 5 served as human endpoints. A mouse was defined as a non-survivor when human endpoints applied and was subsquently sacrificed by cervical dislocation. Mice which did not display any signs of endotoxic shock such as weight or temperature loss and no increasing severeness in score were excluded from the analysis.

### Peripheral blood cytokine concentrations in endotoxic shock

To determine the blood cytokine levels in the endotoxic shock model, mice were injected with corresponding LD_50_ of LPS at six 4-hour spaced time points (n=14). 2h post LPS injection mice were terminally bled by cardiac puncture using a 23G x 1” cannula (Henke-Sass Wolf). Syringes (1ml, Braun) were coated with heparin (Ratiopharm) to avoid blood coagulation. After isolation blood was kept at 4°C for subsequent plasma preparation. To this end, blood was centrifugated for 15min, 370g at 4°C. Plasma was isolated, aliquoted and frozen at -80°C for further analysis.

### Isolation of peritoneal macrophages (PM)

Mice were sacrificed by cervical dislocation. Peritoneal cavity cells (PEC) were isolated by peritoneal lavage with ice cold PBS. Lavage fluid visibly containing red blood cells was dismissed. For RNA measurements or bioluminescence recordings peritoneal macrophages were further purified by MACS sorting (Miltenyi) according to manufacturer’s protocol using mouse/human CD11b Microbeads and LS columns. Eluates containg positively sorted macrophages were analyzed for sorting efficiency by FACS. All steps were performed at 4°C.

### Bioluminescence recordings

PER::LUC protein bioluminescence recordings were used to characterize circadian clock function in peritoneal macrophages, SCN and lung tissue. Mice were sacrificed by cervical dislocation. Brains and lungs were isolated and transferred to chilled Hank’s buffered saline solution, pH 7.2. (HBSS). For tissue culture, 300*µ*m coronal sections of the brain and 500*µ*m sections of the lung were obtained using a tissue chopper. The lung and SCN slices were cultured individually on a Millicell membrane (Millipore) in a Petri dish in supplemented DMEM containing 1*µ*M luciferin (Promega). PECs were isolated by peritoneal lavage and immediately CD11b-MACS sorted. The CD11b-sorted peritoneal macrophages were cultured in Petri dishes in supplemented DMEM containing 1*µ*M luciferin (Promega). For bioluminescence recording, tissues/primary cell cultures were placed in light-tight boxes (Technische Werkstaetten Charite, Berlin, Germany) equipped with photo-multiplier tubes (Hamamatsu, Japan) at standard cell culture conditions. Bioluminescence was recorded in 5min bins. On fifth day in culture PMs were treated with 1*µ*M dexamethason for 1h, followed by a medium change to supplemented DMEM containing 1*µ*M luciferin (Promega). Data were further processed and analyzed using Chronostar 3.0 [49].

### Statistical data analysis

Statistical analysis was performed in GraphPad Prism 8 and R. Normality was tested using Shapiro-Wilk normality test. When data were normally distributed and two parameters were compared, One-way ANOVA with Dunnett’s multiple comparison as a post hoc test was applied. When comparing more than two groups and two parameters a two-way ANOVA with Bonferonni’s post hoc test was applied. Two sample comparison was performed with 2-sided students t-test for normally distributed data and Mann-Whitney U test for non-normally distributed data. Time-of-day dependent mortality experiments: Mortality data were transformed to probability of death (between 0-1) in order to compute sine fit using alogistic regression. The confidence interval was derived from sine fit estimation. After statistical analysis, the mortality data was transformed back to the initial percentages. Cross-correlation analysis of morality and cytokine data: To correlate mortality rates and cytokine levels across all animal models, a permutation of sine fit of the mortality data and plasma cytokine data was used. This boot-strapping (randomization) procedure gives rise to the empirical distribution of correlations. The p-value is the fraction of randomizations that gave a correlation with the opposite denominator, meaning that a p=0.05 means that only 5% of correlations crossed zero correlation threshold. For linear regression analysis of the summary of cytokine and mortality data, the estimated mortality rate determined by sine fit of mortality data was correlated to mean value of plasma cytokine concentration using Spearman’s rank correlation. Bioluminescence recordings were analysed using in-house written software ChronoStar 3.0 [49]. In brief, raw bioluminescence counts were transformed to log-space and trends removed by subtracting the 24h running average. Circadian rhythm parameters were estimated by fitting a damped sine wave to these data. Finally, data were reversely transformed into linear space. Circadian rhythmicity of cytokine time-series was tested using our in-house written software ChronoLyse. In brief, a 24h sine wave was fitted the beforehand log-transformed data and parameters of fit were used to estimate amplitude, phase and mean levels. Rhythmicity was tested by testing against a flat line using F-test.

## Acknowledgments

This work was supported by the Deutsche Forschungsgemeinschaft (DFG, Grants AN 1553-2/1, MA 5108/1-1, HE2168/11-1, SPP 2041).

