## Supplemental data for "Susceptibility rhythm to bacterial endotoxin in myeloid clock-knockout mice"

#### Supplemental Methods

##### Animals

All procedures were authorized by and performed in accordance with the guidelines and regulations of the German animal protection law (Deutsches Tierschutzgesetz). Mice were housed in macrolon type II cages supplied with nesting material, food and water ad libitum at a 12h:12h light/dark (LD) cycle. For endotoxic shock and running wheel experiments mice were individually housed. For all other experiment mice were group-housed. Manipulations during the dark phase of the cycle were performed under infrared light. Male C57Bl/6 mice (Jackson Laboratories strain) mice were purchased from our animal facility (Charité FEM, Berlin, Germany) at 8-10 weeks of age. Homozygous, male *LysM*$^{Cre/Cre}$ (LysM-Cre) [1], myBmal-KO and myClock-KO were bred and raised in our animal facility (FEM, Berlin Germany) and used at 8-12 weeks. Female *LysM*$^{Cre/Cre}$ *Per2:Luc*, either wild-type or homozygous for *Clock*-flox, were bred and raised in our animal facility (FEM, Berlin Germany) and used at 14 weeks.

##### Locomotor activity recording

Male homozygous myClock-KO mice (control mice: LysM-Cre and *Clock*-flox mice, 8-10 weeks of age were individually housed with ad libitum acess to a running wheel in Macrolon typ III cages. Running wheel activity was recorded for 2 weeks in 12h:12h light:dark (LD) followed by 2 weeks of constant darkness (DD). Locomotor activity was recorded and evaluated with ClockLab Analysis (ActiMetrix). Circadian Period (tau) and overall activity analysis was performed in DD phase.

##### Intraperitoneal LPS injection

*E. coli* LPS (O55.B5, Sigma Aldrich) stock solution (10mg/ml) was diluted to appropriate concentration in sterile PBS and thoroughly vortexed before use. LPS injection was performed *i.p.* using Legato 100 Syringe Pump (KD Scientific). The following settings were applied: Mode: infuse only; syringe: BD, plastic, 5ml; rate: 2ml/min. A cannula (26G x 3/8") was attached to Legato 100 by microbore extension line (60cm, MedEx, Smiths Medical) for the LPS injections.

##### Isolation of bone marrow cells

Mice were sacrificed by cervical dislocation. One tibia and femur were excised per mouse. Femur and tibia were flushed with supplemented RPMI 1640 and erythrocytes were lysed using GEYS solution (2min at 4°C). After erythrocyte lysis, cells were suspended in supplemented RPMI 1640 medium and filtered through a 30$\mu$M filter (Miltenyi). Cell numbers were determined using a Neubauer chamber. All steps were performed at 4°C. All centrifugation steps were performed at 4°C, 300g, 7min.

##### Isolation of spleen cells

Mice were sacrificed by cervical dislocation. Spleen was removed and a single cell single cell suspension was obtained using gentleMACS, Miltenyi (program: m_spleen_01) with C-tubes (Miltenyi) in PBS. Next single cell solution was filtered using 100$\mu$M cell strainers (Thermo Fischer). GEYS solution was used for erythrocyte lysis for 2min at 4°C. After erythrocyte lysis cells were suspended in supplemented RPMI 1640 and filtered through a 30$\mu$M filter (Miltenyi). Cell number was determined using a Neubauer chamber. All steps were performed at 4°C. All centrifugation steps were performed at 4°C, 300g, 7min.

#### Generation of whole cell protein lysates

Liver and peritoneal cavity cells were homogenized in ice cold RIPA buffer containing 1x protease-inhibitor-cocktail (Sigma Aldrich) and incubated on ice for 30min. Homogenized cells were then centrifuged at maximum speed for 30min at 4°C to pellet the insoluble cell debris. The supernatant fraction was then removed and used for cellular protein analysis or frozen at -80°C. Protein concentrations were determined using standard BCA assay.

#### Immunoblotting

Samples were denaturated for SDS-PAGE in NuPAGE SDS Sample Buffer (4x) (Invitrogen) containing 0.8% 2-$\beta$-mercaptoethanol (Sigma) and boiled for 5-10min at 95°C. SDS-PAGE using 4-12% Bis-Tris gels (Thermo Scientific) at 200V for 60min in NuPAGE MES SDS Running Buffer. Proteins were transferred to a nitrocellulose membrane (0.45$\mu$m) using a tank transfer system (wet transfer). NuPAGE transfer buffer, containing 20% Methanol, was cooled with an ice block to prevent overheating during the transfer. The transfer was run for 120min at 90V. Following the transfer, the membrane was blocked in TBS-T with 5% non-fat, dry milk for 1-2h at RT. After a washing step in TBS-T (3 x 10min), the membrane was placed in the primary antibody solution (TBS-T with 5% non-fat, dry milk) and gently shaken overnight at 4°C. The membrane was then washed in TBS-T (3 x 10min) and incubated with the HRP-conjugated secondary antibody (Santa Cruz Biotechnologies) in TBST-T for 2h at RT. After another washing step in TBS-T (3 x 10min), a chemiluminescence reaction was performed with Super SignalWest Pico substrate (Pierce). The protein bands were visualized using the ChemoCam detection system (Intas). The following primary antibodies were used: murine CLOCK - rabbit anti-mCLock (Bethyl Laboratories, A302-618A) , murine BMAL1 - rabbit anti-mBMAL1 (kind gift from Micheal Brunner) , murine ACTINB - mouse anti mBactin (Sigma, A5441). Secondary antibodies used: goat anti-mIgG-HRP (SantaCrz Biotechnology, sc-2005), donkey anti-rbIgG-HRP (SantaCrz Biotechnology, sc-2005).

#### Single- and multiplex immunoassays

Singleplex assay: murine IL-6 plasma concentration was determined by ELISA according to manufacturer's protocol (Ebioscience) in a 96-well format (Corning). Plasma samples were diluted 1:200 in supplied assay buffer. Absorption was measured at 470nm. Reference wavelength was measured at 560nm by Infinite F200Pro plate reader (Tecan). Multiplex assay: 13 cytokines (CCL2/MCP-1, CCL3/MIP-1$\alpha$, CCL4/MIP-1$\beta$, CCL7/MCP-3, CXCL5, IL-1$\alpha$, IL-1$\beta$, IL-10, IL-12p40, IL-18, Eotaxin, Rantes/CCL5, TNF$\alpha$) were assessed using the ProCartaPlex, Mix and Match, Mouse 13-Plex (Affymetrix, eBioscience). ProCartaPlex was performed as described in manufacturer's protocol in a 96 well format (eBioscience). All washing steps were performed using a hand held magnetic washer (eBioscience). Data were acquired using a MagPix (Luminex) detection device. Data evaluation was performed using ProcartaPlex Analyst v.1.0 (eBiosciences).

#### Isolation and quantification of RNA

Total RNA was isolated using the PureLink RNA Mini Kit (Ambion) according to the manufacturer's manual. In addition, an on-column DNA digestion was performed using PureLink DNase Set (Life Technologies). RNA was quantified by measuring the absorption at 260nm with NanoDrop 2000C (Thermo Scientific).

#### Quantitative real-time PCR

Total RNA was reverse-transcribed to cDNA using random hexamers to prime reverse transcriptase reaction. cDNA was diluted 1:10 in $H_2O$ for use in qRT-PCR. qRT-PCR was performed using a 2-step protocol with the following primer-sets: primer-sets for mCry1, mCry2, mDpb, mNr1d1, mNpas2, mPer1, mPer2 were purchased from Qiagen (QT00117012, QT00168868, QT00103089, QT00164556, QT00108647, QT00113337, QT00198366, respectively). mGapdh, fwd: ACGGGAAGCTCACTGGCATGGCCTT, rev: CATGAGGTCCACCACCCTGTTGCTG; mBmal1 primers were designed to characterize myBmal-KO mice. Forward primer (fwd: GGACACAGACAAAGATGACCC) binds upstream of exon 8 and the reverse primer (rev: TTTTGTC-

CCGACGCCTCTTT) within exon 8 of *Bmal1*. Thus after successful Cre recombination, exon 8 is deleted and no PCR product is detectable. *Clock* primers were designed to characterize myClock-KO mice. Primers bind in exon 5 (fwd: ATTGGTGGAAGAAGATGACAAGGA) and in exon 6 (rev: TACCAGGAAGCATAGACCCC) of clock. As exon 5 and 6 are flanked by loxP sites, after successful Cre recombination no PCR product is amplified.

##### Flow cytometry (FACS)

Two panels of antibodies were established to target a broad range of immune cells in various sites of the organism. Before each experiment, antibody mix of both panels were prepared and kept on 4°C for labeling of all samples of the respective experiment in order to minimize intra-experimental variability. All antibody-mixes were prepared in FACS buffer containing 1:50 FcR blocking reagent. FACS staining of samples: 100$\mu$L of the cell suspensions were transferred into a 96-well plate and spun down for 7min, 300g at 4°C. Supernatant was carefully discarded and pellet re-suspended in 50$\mu$L of master-mix containing one of two antibody panels. Cells were incubated for at least 30mins at 4°C in darkness. Subsequently 200$\mu$L FACS buffer was added and cells were centrifuged for 7min, 300g at 4°C. Cells were washed twice using 200$\mu$L FACS buffer each before fixation in 200$\mu$L 4% PFA for 30min at RT. Finally, cells were spun down and re-suspended in 200$\mu$L FACS buffer and stored at 4°C for up to a week before FACS data acquisition in a FACS CantoII (BD Biosciences).

#### Supplemental Tables

##### Supplemental Table 1

| Score | Behavior, Phenotype |
|---|---|
| 0 | normal, no behavioural abnormalities |
| 1 | slightly decreased speed of course of movement |
| 2 | slightly lethargic |
| 3 | “slow motion" movements, ruffled fur, hunched posture |
| 4 | lethargic, movements are not specific to cue, strong bar-grip-reflex |
| 5 | completely lethargy, body position is not self-determined, decreased bar-grip-reflex |

**Supplemental Table 1.** Scoring system to determine human endpoint in endotoxic shock experiments.

### Supplemental Figures

#### Supplemental Figure 1

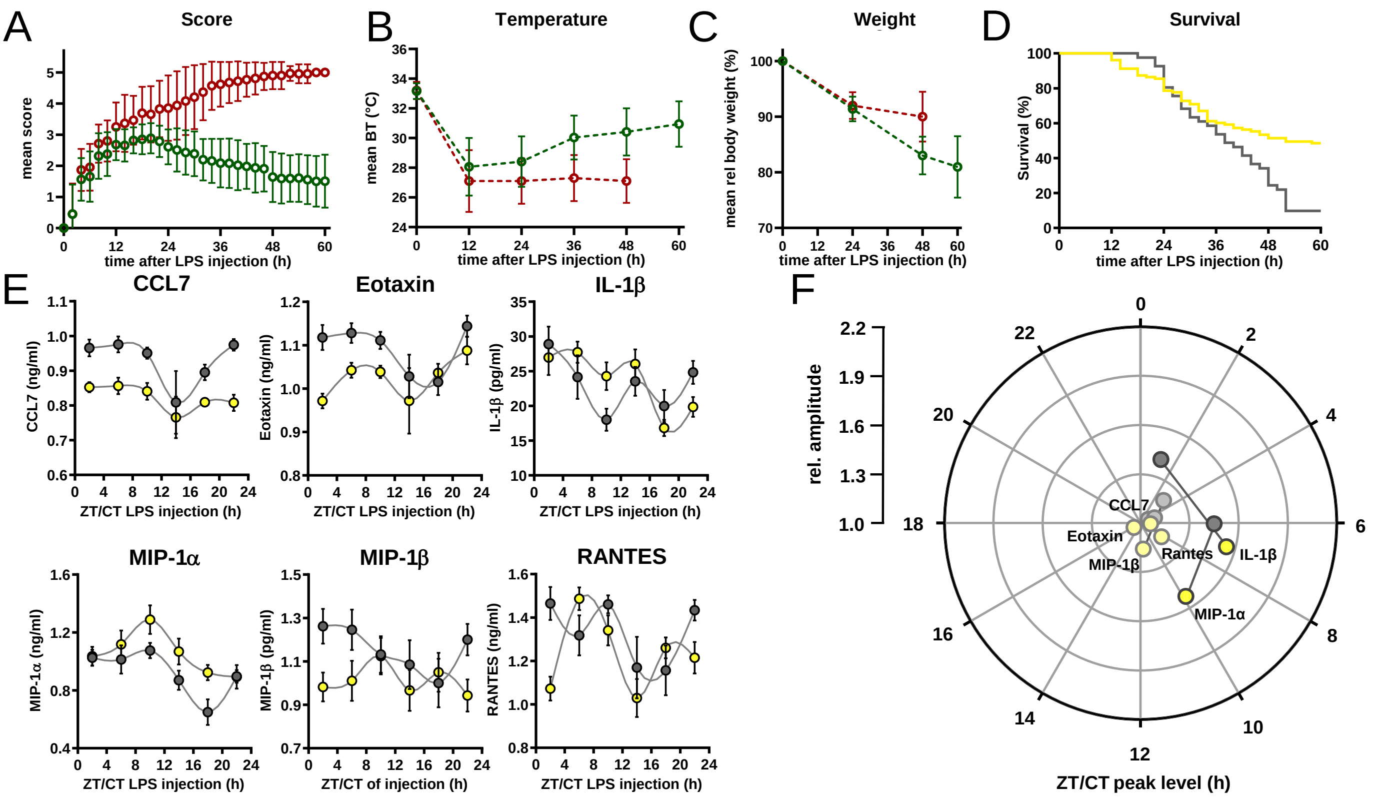


**Supplemental Figure 1. A-C)** Score, body temperature and weight development of C57Bl/6 mice kept in LD and challenged with LPS (30mg/kg, *i.p.*) **A)** Mean score development, **B)** mean body temperature development and **C)** mean body weight development in survivors (green circles, n=49) and non-survivors (red circles, n=54 at time point 0). Error bars represent SD. **D)** Kaplan-Meier survival curves of C57Bl/6 mice kept either in LD (yellow line) or DD (grey line) and challenged with LPS (30mg/kg, *i.p.*). Data have been aligned to time after challenge with LPS, irrespective of time of day of challenge (same experiment as shown in Fig. 1C). Survival curves differ significantly (p=0.0006). **E)** Time-of-day dependent cytokine profiles (complementing data shown in Fig. 1E) in peripheral blood of C57Bl/6 mice (n=10 per time point) challenged with half-lethal doses of LPS and sacrificed 2 hours later. LPS dosage for mice kept in LD (yellow, 30mg/kg) or DD (grey, 13mg/kg). **F)** Relative amplitudes and phases of cytokines shown in E). Light-colored circles represent non-significant circadian rhythms (p-value>0.05 ) as determined by non-linear least square fit and consecutive F-test (see also Methods section).

#### Supplemental Figure 2

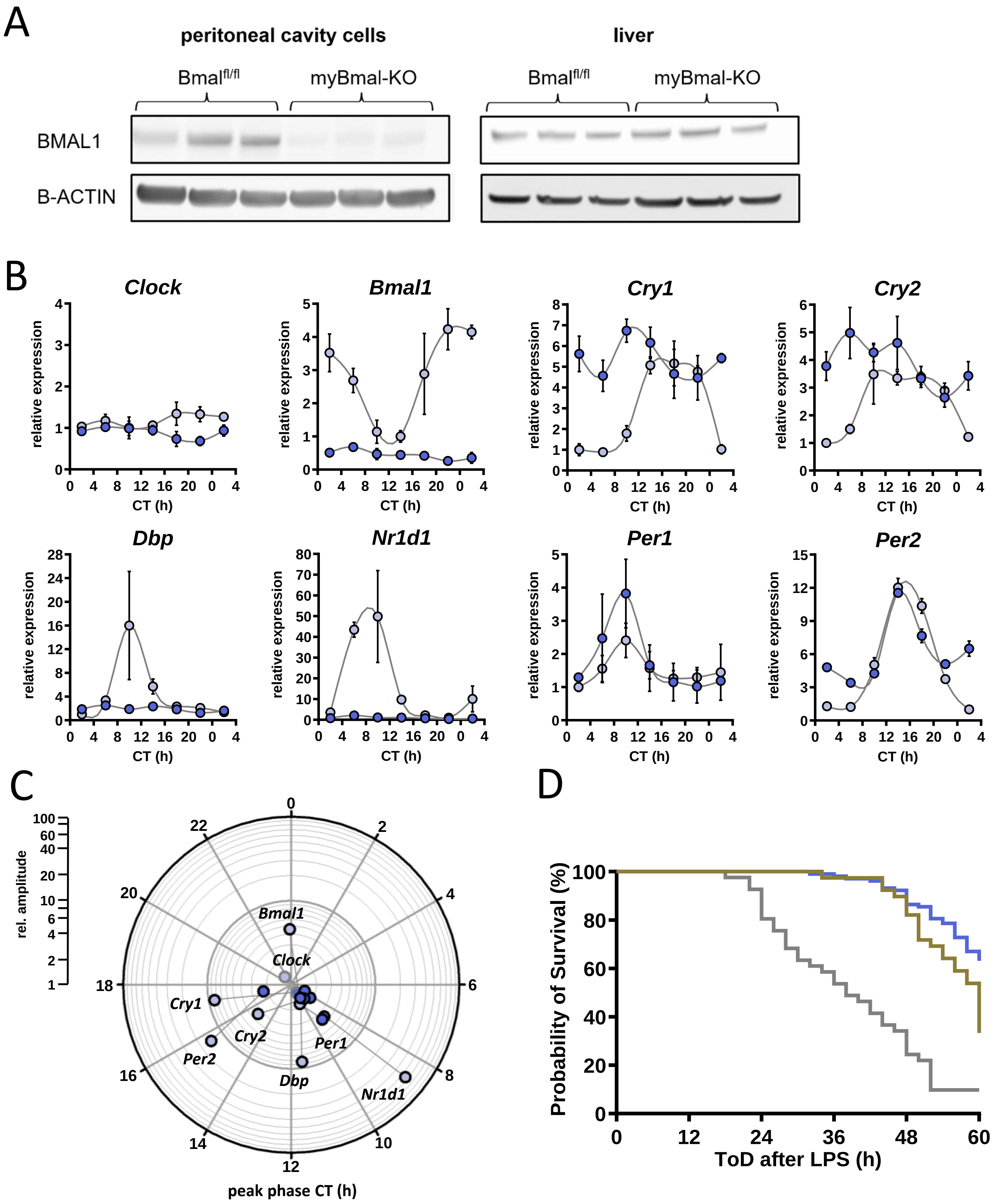


**Supplemental Figure 2. Molecular characterization of myBmal-KO mice. A)** Western blots of whole cell lysates from peritoneal cavity cells or liver of control (Bmal-flox$^{+/+}$) and myBmal-KO mice (n=3). **B)** Relative mRNA levels of selected clock genes in peritoneal macrophages from Bmal-flox$^{+/+}$ (light blue circles) or myBmal-KO (blue circles) mice at indicated circadian times. Phase and amplitude information are depicted in C) as analyzed by Chronolyse. Non-significant circadian expression ($p>0.05$) are indicated by grey shaded circles for both myBmal-KO and LysM-cre control. **C)** Kaplan-Meier survival curve of mice kept in DD and challenged with LPS (30mg/kg, *i.p.*). Grey line refers to C57Bl/6 mice (wild-type control), brown line to LysM-cre control and purple line to myBmal-KO. Data shown have been aligned to time after challenge with LPS, irrespective of time of day of challenge (same experiment as shown in Fig. 1C and 2B, respectively). Survival curves of wild-type and myBmal-KO as well as LysM-cre$^{+/+}$ and myBmal-KO differ significantly ($p<0.0001$ and $p=0.0031$, respectively, log-rank tests).

#### Supplemental Figure 3

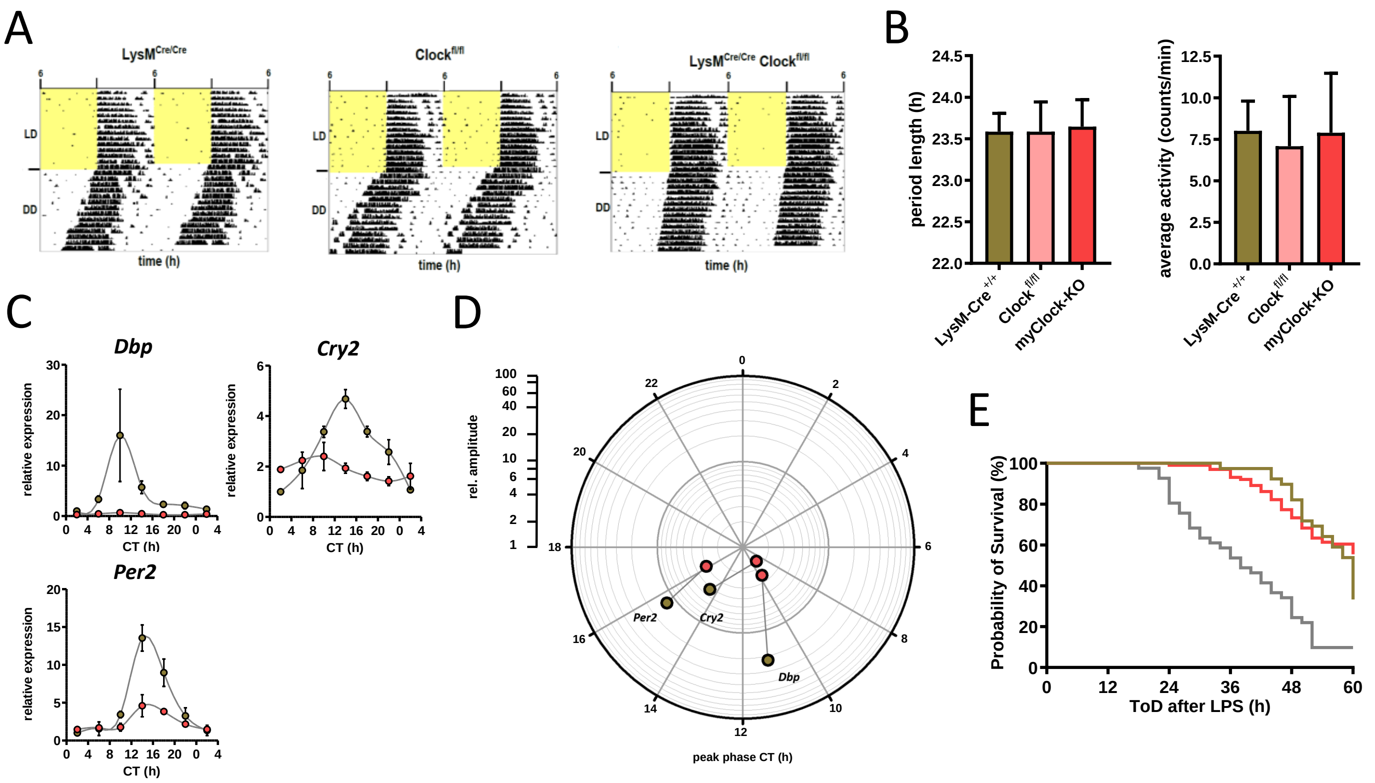


**Supplemental Figure 3. Phenotypic and molecular characterization of myClock-KO mice. A)** Locomotor activity recordings from control (LysM-cre$^{+/+}$ and Clock-flox$^{+/+}$) and myClock-KO mice. Representative examples are shown and quantified (n=7-13) in **B)** for circadian periods as well as overall activity. **C)** Relative mRNA levels of selected clock genes (complementing data of Fig. 3C) in peritoneal macrophages from LysM-cre$^{+/+}$ (brown circles) or myClock-KO (red circles) mice at indicated circadian times. Phase and amplitude information are depicted in **D)** as analyzed by Chronolyse. Non-significant circadian expressions ($p > 0.05$) are indicated by light shaded circles for both myClock-KO and LysM-cre control. **E)** Kaplan-Meier survival curve of mice kept in DD and challenged with LPS (30mg/kg, *i.p.*). Grey line refers to C57Bl/6 mice (wild-type control), brown line to LysM-cre control and red line to myClock-KO mice. Data have been aligned to time after challenge with LPS, irrespective of time of day of challenge (data are from experiments shown in Fig. 1C and 3F/G, respectively). Survival curves of wild-type and myClock-KO differ significantly ($p < 0.0001$, log-rank test).

#### Supplemental Figure 4

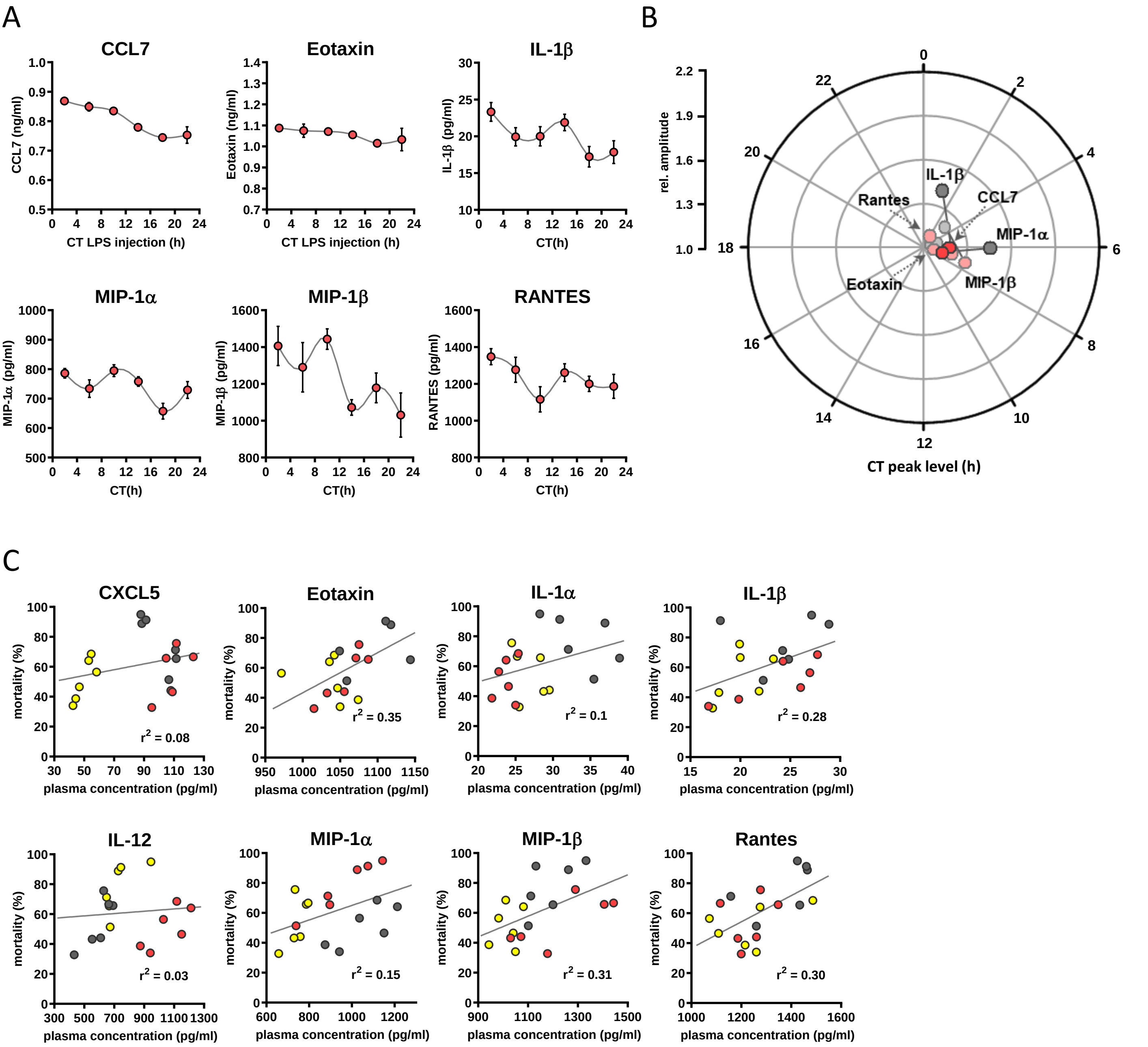


**Supplemental Figure 4. Circadian time dependent cytokine levels in plasma of myClock-KO mice complementing data in figure 4. A)** Plasma cytokine levels in myClock-KO mice kept in constant darkness, two hours after *i.p.* injection of 30mg/kg LPS. Data represent mean ± SEM (n=14 per time point). **B)** Polar plot showing amplitude and phase distribution of cytokines from A), red circles, myClock-KO and wild-type DD, grey circles (Fig. 1E). Light circles indicate non-significant circadian abundance (p-values>0.05, non-linear least square fit statistics by Chronolyse). **C)** Overall correlation of cytokine levels with mortality independent of time-of-day of LPS injection and mouse model. Colors indicate data source (wild-type, LD - yellow; wild-type, DD - gray; myClock-KO, DD - red; linear regression - gray line; statistics: spearman correlation).

#### Supplemental Figure 5A

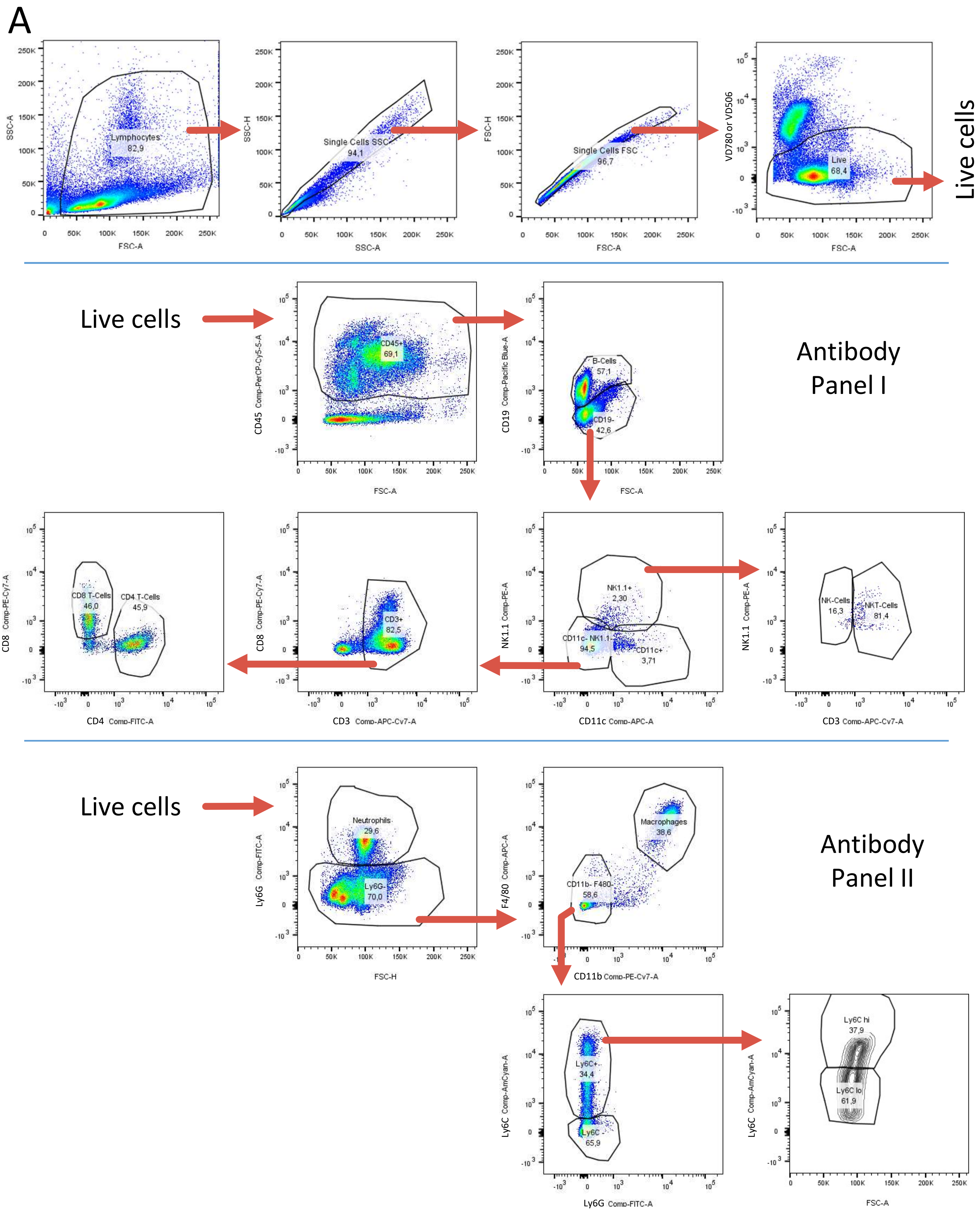

#### Supplemental Figure 5B

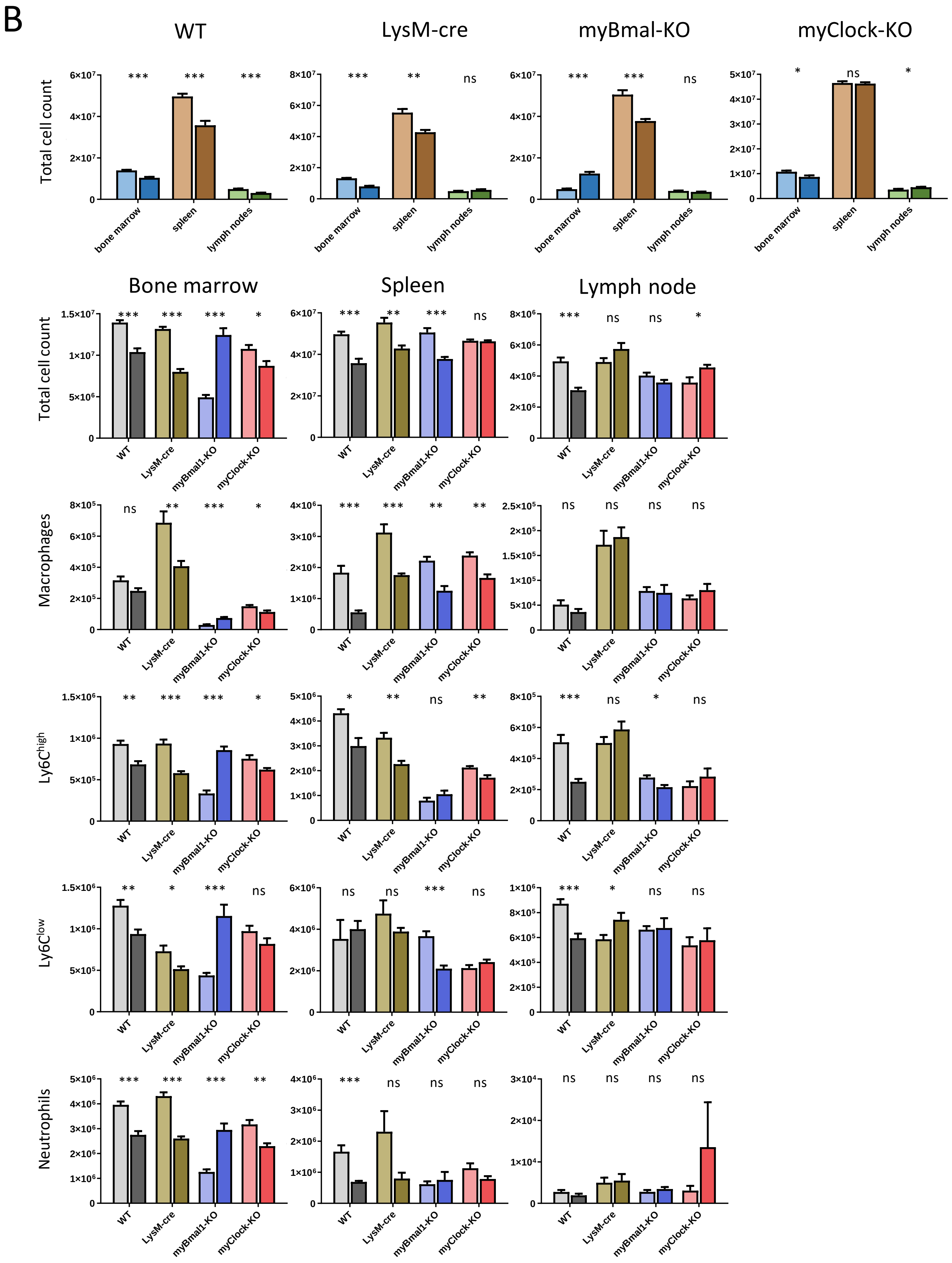

#### Supplemental Figure 5C

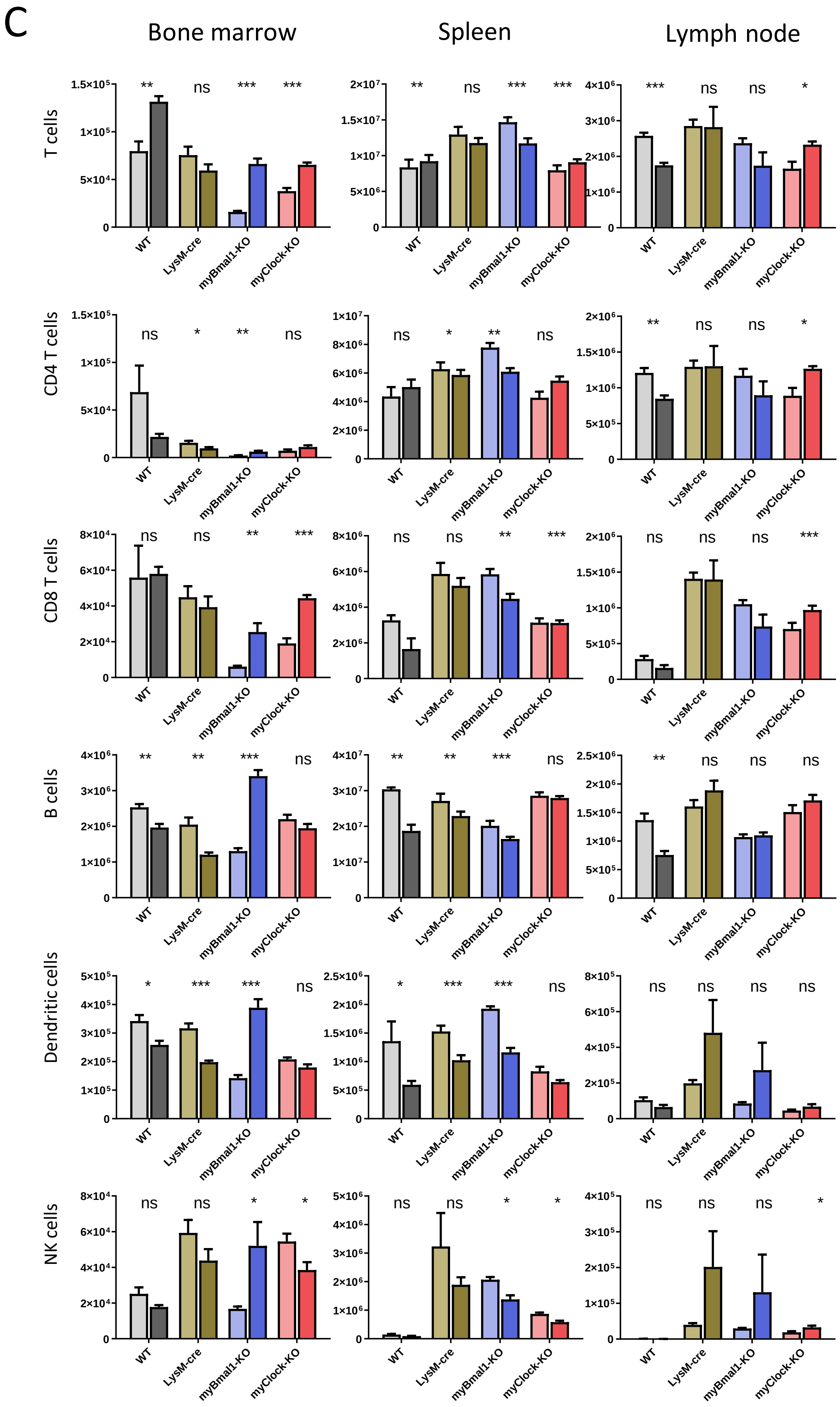

#### Supplemental Figure 5D

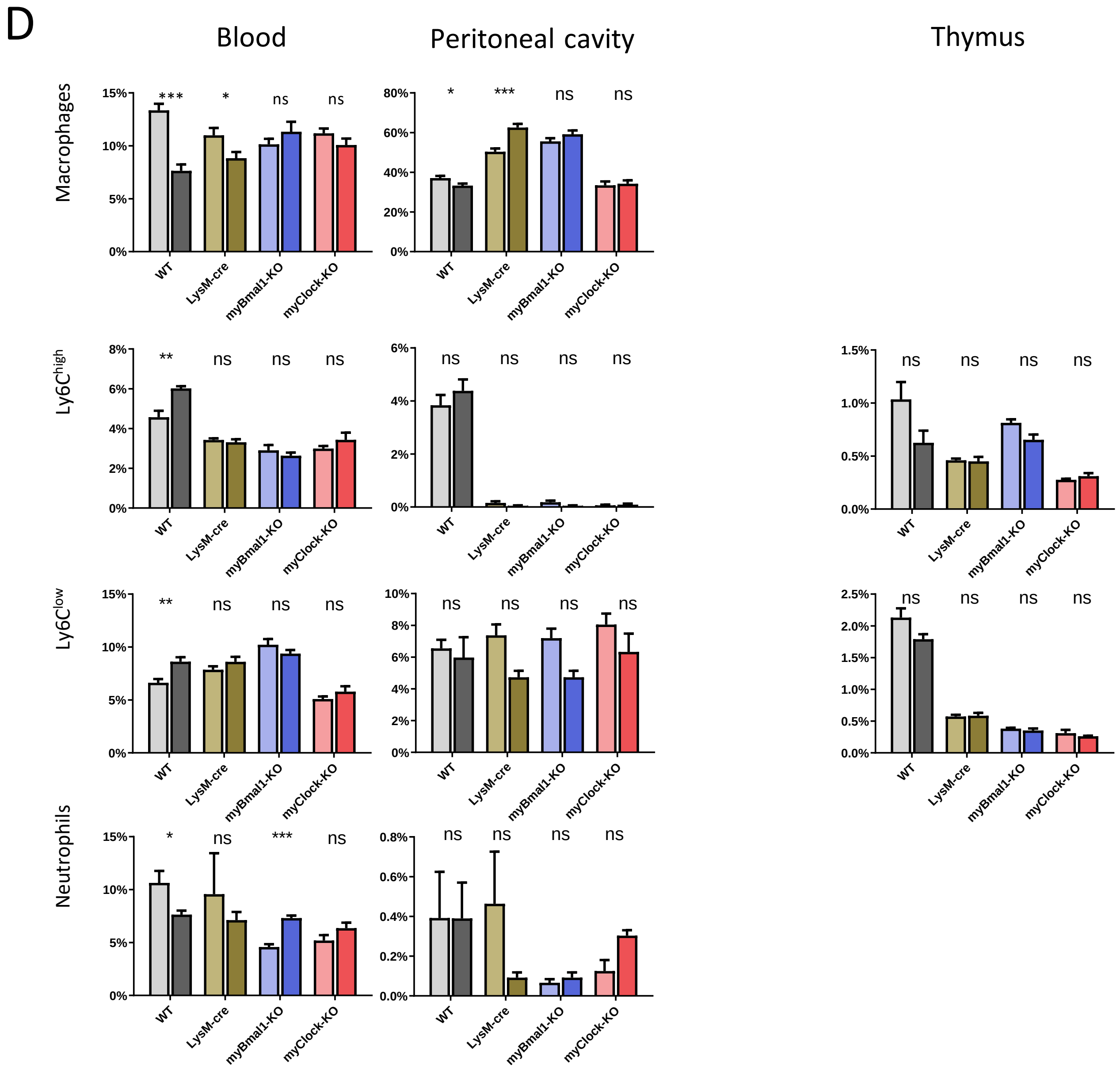

#### Supplemental Figure 5E

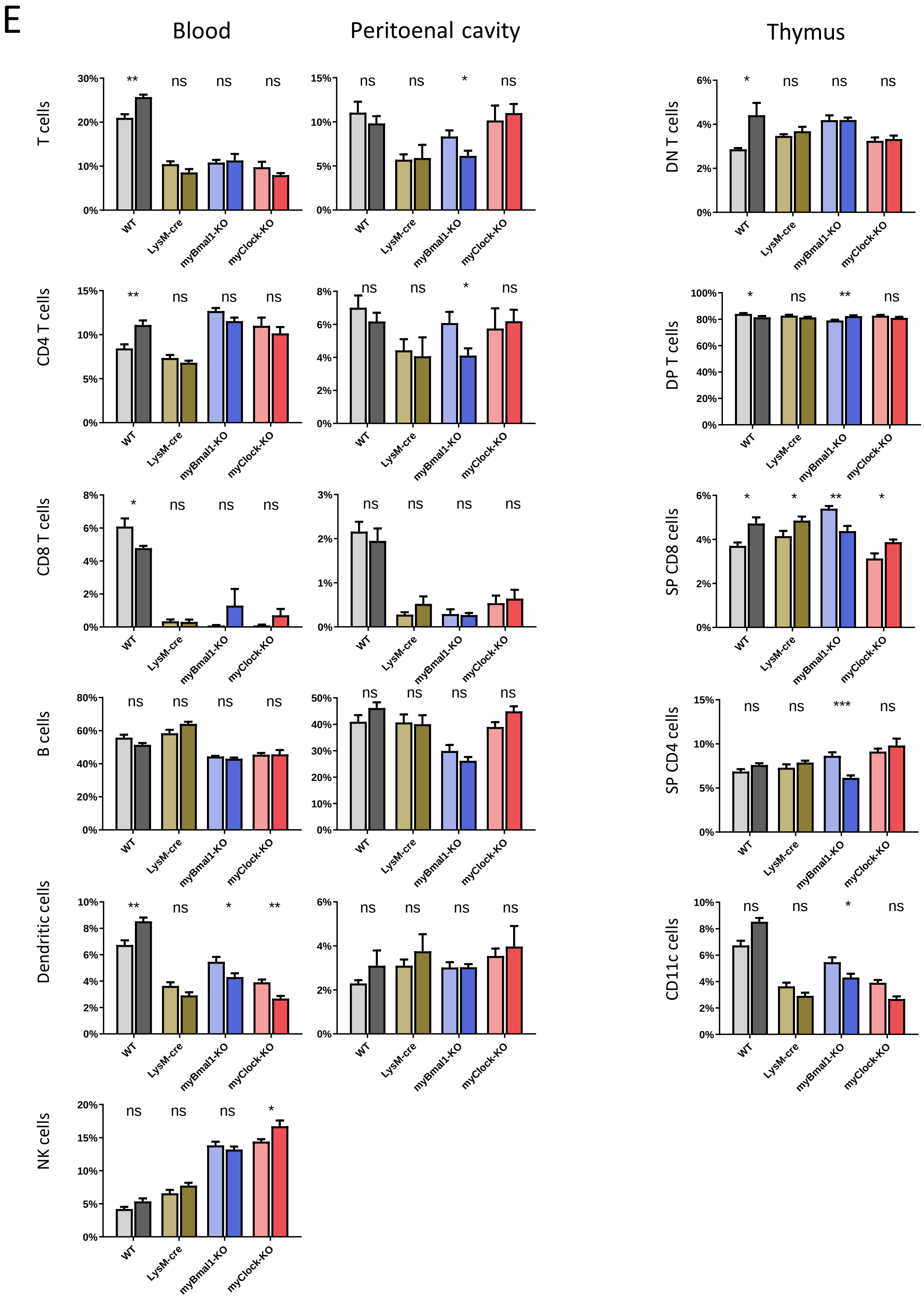

#### Supplemental Figure 5 Legend

**Supplemental Figure 5. A)** FACS gating strategy for identification of lymphoid and myeloid immune cell subtypes for quantifications shown in Fig. 5 as well as in Suppl. Fig. 5B-E. **B)** Total cell counts as well as cell counts for indicated immune cell subtypes derived from antibody panel II (Suppl. Fig. 5A) in bone marrow, spleen and lymph nodes of control as well as myeloid clock knockout mice at CT8 and CT20. Colors of bars refer to scheme of experimental setup shown in Fig. 5A. Asterisks indicate level of significance as determined by student t-test. **C)** Cell counts for indicated immune cell subtypes derived from antibody panel I (Suppl. Fig. 5A) in bone marrow, spleen and lymph nodes of control as well as myeloid clock knockout mice at CT8 and CT20. **D)** Frequencies of immune cell subtypes derived from antibody panel II (Suppl. Fig. 5A) in blood, peritoneal cavity and thymus of control as well as myeloid clock knockout mice at CT8 and CT20. **E)** Frequencies of immune cell subtypes derived from antibody panel I (Suppl. Fig. 5A) in blood, peritoneal cavity and thymus of control as well as myeloid clock knockout mice at CT8 and CT20. Significance levels in all graphs: ns $p>0.05$, * $p<0.05$, ** $p<0.01$, *** $p<0.001$.
